# Recent divergence and microgeographic genetic structure in an endangered Australian songbird: the southern black-throated finch

**DOI:** 10.1101/2025.04.17.649380

**Authors:** Daniel M. Hooper, Kelsie A. Lopez, Bronwyn G. Butcher, Irby J. Lovette, Simon C. Griffith

## Abstract

Anthropogenic habitat loss and climate change threaten global biodiversity. Effective conservation management requires a detailed understanding of geographic structure, genetic diversity, and demography of threatened species. The black-throated finch, *Poephila cincta*, is an Australian songbird with two subspecies: *atropygialis* and *cincta*. The southern subspecies, *cincta*, has experienced an ∼80% range contraction over the last century and is listed as endangered but genetic surveys of it are incomplete. Here, we use a combination of reduced representation and whole genome sequencing to examine genetic differentiation, spatial genetic structure, and demographic history in both forms of this species. We find that *atropygialis* and *cincta* are genetically distinct despite a history of divergence with gene flow and geographically isolated by a biogeographic barrier known as the Einasleigh Uplands. Since they last shared a common ancestor ∼400,000 years ago, the two subspecies have experienced distinct demographic trajectories: population expansion in *atropygialis* and population decline in *cincta*. We find that the two remnant population centers of *cincta*, from the Galilee Basin and the Townsville Coastal Plain, each represent genetically distinct lineages that last shared appreciable levels of gene flow ∼4,000 years ago. Moreover, we report striking microgeographic genetic structure from the Townsville Coastal Plain between populations <20 km apart associated with barriers to dispersal caused by anthropogenic habitat modification over the last 50 years: namely the construction of the Ross River Dam. Our findings highlight the urgent need for a conservation approach that prioritizes habitat restoration to re-establish population connectivity in the endangered southern black-throated finch.

## INTRODUCTION

Habitat loss, degradation, and fragmentation constitute some of the gravest modern risks to biodiversity (Brooks et al., 2002; Hanski 2011; Groom et al., 2012; Diaz et al., 2019). A species’ extinction risk is elevated through such landscape alterations largely by lowering connectivity between populations. A central conceit of the conservation genetics paradigm holds that small and isolated populations are especially susceptible to the loss of genetic diversity through processes such as random genetic drift and inbreeding (Hedrick 2001; Frankham 2005; Ouborg et al., 2006). Much like a spiraling feedback loop, shrinking populations with declining genetic diversity are at greater risk of extinction due to the increased frequency of deleterious alleles that can result in inbreeding depression (Charlesworth and Charlesworth 1987; Charlesworth 2009), an elevated susceptibility to disease (Spielman et al., 2004; King and Lively 2012), and a heightened vulnerability to stochastic environmental change (Lande 1993; Reed 2004). Moreover, the loss of genetic variation within populations can critically hamper the resilience and adaptive capacity of a species to further environmental change (Reed and Frankham 2003; Hoffmann and Sgrò 2011; Urban 2015; Exposito-Alonso et al., 2022).

Australia has one of the highest rates of continental biodiversity loss worldwide, driven largely by habitat destruction, invasive species, and climate change (Laurance et al., 2011; Woinarski et al., 2015; Ward et al., 2019). Granivorous bird species have been disproportionately affected, as they depend on intact fire-adapted grassland ecosystems for food and nesting sites, making them highly susceptible to habitat fragmentation and degradation (Franklin 1999; Franklin et al., 2005). Approximately one-quarter of all granivorous bird species show evidence of population decline in northern Australia over the last 150 years (Franklin 1999), several taxa are listed as endangered, and two are extinct (Garnett and Baker, 2021).

The black-throated finch (*Poephila cincta*) is a songbird endemic to the northeastern tropics of Australia and comprises two subspecies that differ in rump color: black in the northern subspecies *atropygialis* and white in the southern subspecies *cincta* (Higgins 2006). The range of *atropygialis* extends across much of the Cape York Peninsula west to the Flinders River and south to the Atherton Tablelands in tropical northern Queensland. Subspecies *cincta* historically ranged from the Atherton Tablelands south to northeastern New South Wales (Figure 1A). Like many granivorous Australian bird species, however, *cincta* has experienced a severe and well-documented range contraction by ∼80% in the last century due to widespread habitat conversion to agriculture, habitat degradation due to high intensity grazing, and changes to historical fire management regimes (Franklin 1999; Franklin et al., 2005; Reside et al., 2019). The range of *cincta* has decreased to such an extent that it was listed as endangered in 2005 (Environment Protection and Biodiversity Conservation Act, 1999), declared ‘presumed extinct’ in New South Wales in 2016 (Biodiversity Conservation Act, 2016), and is now likely to persist in large numbers in only two locations of its former range: the Townsville Coastal Plain and the Galilee Basin (Vanderduys et al., 2016; Mula-Laguna et al., 2019). Despite these official listings, habitat loss has continued in recent years and the Galilee Basin population is currently threatened by the development of the Carmichael Coal Mine and Rail Project and other related mining projects (Reside et al., 2019). In contrast, the current range of northern subspecies *atropygialis* appears to be more stable and exhibits no documented signs of decline (Franklin 1999; Garnett and Baker, 2021). While hybridization between *atropygialis* and *cincta* has been reported in the historical literature (Keast 1958), the extant distributions of these taxa would appear to preclude the possibility of contact between them today. The only assessment to date of genetic differentiation between black-throated finch subspecies found no evidence of contemporary admixture but was limited by its sparse geographic sampling and reliance on a relatively small panel of microsatellite markers (Tang et al., 2016). Twin goals of this study, therefore, are to develop a clearer understanding of genomic divergence between black-throated finch subspecies and to explore their respective demographic histories.

**FIGURE 1.**
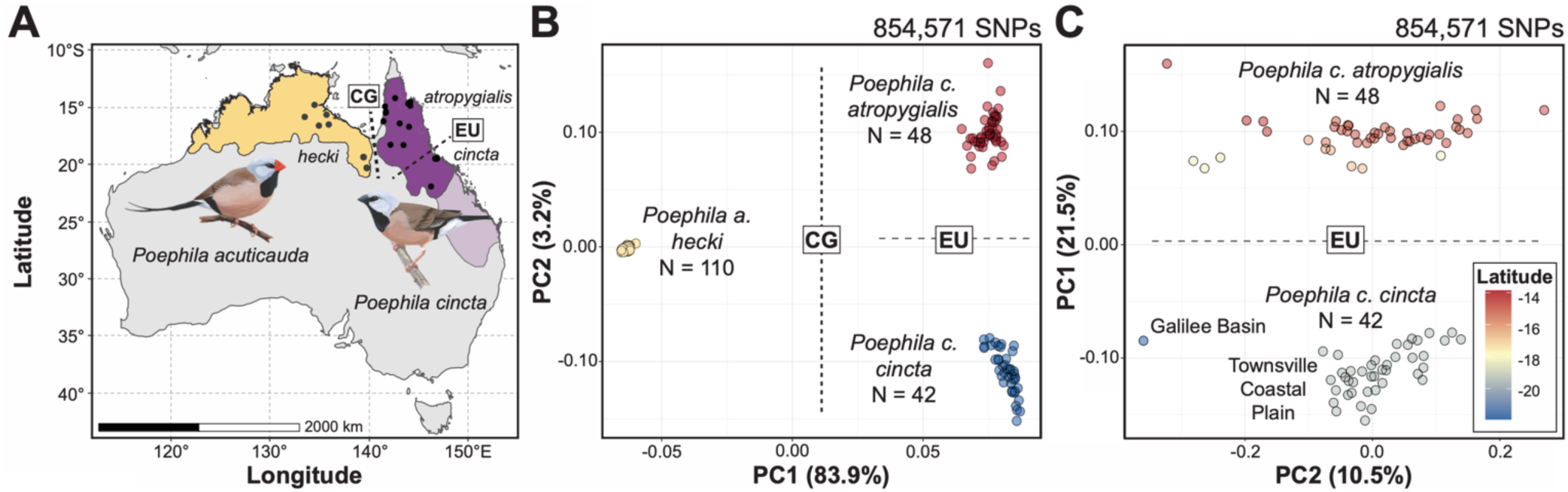
Geographic distribution and genomic divergence between both black-throated finch (*Poephila cincta cincta* and *P. c. atropygialis*) subspecies and long-tailed finch (*Poephila acuticauda hecki*). (A) Estimated range distribution of long-tailed finch (yellow) and black-throated finch (purple). Recently extirpated regions of the black-throated finch geographic range (subspecies *cincta*) are represented with a lighter shade of purple. The geographic locations of populations sampled in this study are represented as black circles (see Table S1 for precise locations). The dashed lines represent the approximate geographic locations of two relevant biogeographic barriers: the Carpentarian Gap (CG) and the Einasleigh Uplands (EU). (B) Principal components analysis based on genome-wide SNP variation using all WGS dataset samples from both species (N = 200) and (C) only samples from the black-throated finch (N = 90). Biogeographic barriers CG and EU are depicted as labeled dashed lines separating samples from either side of them. Samples are color-coded by taxon in (B) and by the latitude (i.e., LAT) of the population they were sourced from in (C).

Our approach in this study has three components. First, we leverage genomic resequencing data to ascertain the extent of genetic divergence and spatial genetic structure between black-throated finch subspecies and its closest relative, the long-tailed finch (*P. acuticauda*). We focus particularly on evaluating spatial genetic structure within and between the two remaining large populations of subspecies *cincta* located in the Townsville Coastal Plain and the Galilee Basin, respectively. Second, we examine the extent to which genetic diversity and evidence of inbreeding vary between populations of each species. We especially seek to evaluate how range fragmentation over the last century has affected diversity in *cincta*. Third, we use demographic reconstruction analyses to infer the respective divergence times, effective population sizes, and migration patterns between black-throated finch subspecies. Taken together, we hope that this survey of genomic differentiation and genetic structure will inform and motivate conservation efforts centered around preventing and reversing habitat loss in the southern black-throated finch (subspecies *cincta*).

## METHODS

### Samples and sequencing

Linked-read and short-read whole genome sequencing (WGS) data comes from previously published work (Hooper et al., 2024) while the reduced-representation sequencing data (ddRAD) was generated for this study, as described below. We examined samples from both black-throated finch subspecies (*Poephila cincta cincta*, hereafter ‘*cincta’*, and *P. c. atropygialis*, hereafter ‘*atropygialis’*) and the eastern subspecies of the long-tailed finch (*P. a. hecki*, hereafter ‘*hecki’*) (Fig. 1A).

Linked-read (LR) whole genome sequencing data was originally generated by Hooper et al. (2024) (NCBI BioProject PRJNA1101033 and PRJNA1147746). In brief, LR genomic libraries were prepared following minor modifications to the 96-plex “haplotagging” protocol from Meier et al. (2021). This dataset included 96 black-throated finch (*cincta*, N = 48; *atropygialis*, N = 48) and 1133 long-tailed finch samples representing both subspecies (*P. a. acuticauda* and *P. a. hecki*) and their hybrids. The full bioinformatic pipeline for processing this data is described in detail in Hooper et al. (2024). We recovered an average of 8.4 million paired end reads per sample and a median depth of coverage of 1.4× for the 1229 samples with LR WGS data (Table S1). Data from all 1229 samples was used through the genotype calling stage, after which only a subset of samples relevant to this project were retained – as described below.

Short-read (SR) whole genome sequencing data was originally generated by Singhal et al. (2015) and Hooper et al. (2024). In brief, we analyzed WGS data from eight long-tailed finch (subspecies *hecki*; Singhal et al., 2015) and two black-throated finch samples (*cincta*, N = 1; *atropygialis*, N = 1; Hooper et al., 2024). WGS data for the long-tailed finch samples was available through ENA (PRJEB10586; Singhal et al., 2015) and was generated on an Illumina HiSeq 2000 (100 bp paired end reads). WGS data for the black-throated finch samples was available through NCBI (PRJNA1101033 and PRJNA1147746) and was generated on an Illumina HiSeq 4000 (150 bp paired end reads). Read data was processed and mapped to the zebra finch reference genome (GCF_003957565.2) as described in Hooper et al. (2024). We recovered an average of 191.2 million paired end reads per sample and a median depth of coverage of 29.4× for the 10 samples with SR WGS data (Table S2).

Reduced representation (ddRAD) sequencing data was prepared as follows. We obtained blood or muscle tissue from a total of 281 black-throated finch samples (*cincta*, N = 164; *atropygialis*, N = 117). All samples came from museum vouchered specimens loaned by the Australian National Wildlife Collection (ANWC; Canberra, Australia). We also included 24 long-tailed finch samples as an outgroup (all of subspecies *hecki*). Genomic DNA was extracted using a DNeasy Blood & Tissue Kit (Qiagen, California, USA) following the manufacturer’s protocol. Double-digested restriction-site associated DNA markers (ddRADtags) were generated following the protocol of Peterson et al. (2012) with modifications established in Thrasher et al. (2018). We used 200-500 ng of DNA at a standardized concentration of 20 ng/uL for each sample. DNA concentrations were quantified using the Qubit dsDNA Broad Range Quantification Assay Kit (Thermo Fisher Scientific, USA) on a Qubit fluorometer (Life Technologies, NY, USA). DNA was digested with *SbfI* and *MspI* (New England BioLabs, MA, USA) and the ends of the digested genomic DNA fragments were ligated using T4 DNA ligase (New England BioLabs) to P1 and P2 adapters (Peterson et al., 2012). We pooled samples with unique P1 barcodes into different indexing groups post digestion/ligation. We cleaned each pooled sample using 1.5× Agencourt AMPure XP beads (Backman Coulter, CA, USA) and fragments between 400 and 900 bp were size selected using Blue Pippin (Sage Science, MA, USA). We performed 11 PCR cycles with Phusion High-Fidelity DNA Polymerase (New England Biolabs) to incorporate the full Illumina TruSeq primer sequences into the library. Finally, we used an additional AMPure clean-up step to remove small fragments before visually evaluating libraries on a fragment Bioanalyzer (Agilent Technologies, CA, USA) to determine fragment size distribution. Indexing groups were combined and sequenced on one Illumina NextSeq 500 lane (150 bp single end reads) at the Cornell University Institute for Biotechnology and two Illumina HiSeq X lanes (150 bp paired end reads) operated by a commercial service provider (Novogene, CA, USA).

We demultiplexed raw sequencing data using the process_radtags function in STACKS v2.59 (Catchen et al., 2011; Catchen et al., 2013). We only kept reads that passed default STACKS quality filters, the Illumina quality filter, contained both an intact SbfI and MspI RAD site, contained one of the unique barcodes with one mismatch allowance, and did not contain Illumina indexing adapters. All reads that passed these quality filters were then trimmed at their 3’ end to the length of the shortest sequence (i.e., 147 bp) using Trimmomatic v0.39 (Bolger et al., 2014). Read data was mapped to the zebra finch reference genome using BWA mem (Li 2013). We recovered an average of 3.5 million reads per sample and a median depth of coverage for the sequenced genome of 25.9× for the 305 samples with ddRAD data (Table S3).

The WGS and ddRAD datasets have broadly overlapping sets of samples. Following data quality filtering, the majority of the black-throated finch samples with WGS data described above were represented in the set of samples with ddRAD data (73 of 90 samples) and all the long-tailed finches with ddRAD data were represented in the set of samples with WGS data (Table S1 and Table S3).

### Variant discovery and genotyping

We generated an initial set of genomic variants found in our set of 1239 samples with WGS data for the 30 largest chromosomes in the zebra finch reference using the mpileup function of bcftools v1.9 (Li 2011; Danecek et al., 2021). We restricted analyses to these 30 chromosomes, together representing 98.3% of the genome, because previous work (Hooper et al., 2024) detected a roughly two-fold reduction in mapping performance and subsequent variant calling for the 10 smallest chromosomes <2.7 Mb in length. A subset of biallelic SNPs were then selected after bcftools filtering for proximity to indels (i.e., -g3) and based on variant quality and sequencing error artefacts (%QUAL<500 || AC<2 %QUAL<50 || %MAX(AD)/%MAX(DP)<=0.3 || RPB<0.1 && %QUAL<50).

This quality-filtered subset of variants was then pruned of sites that overlapped with annotated repetitive regions of the zebra finch genome called by repeatModeler2 (Flynn et al., 2020). A total of 37.59 million SNPs remained after initial quality filtering and repeat masking. Following the approach of Hooper et al. (2024), we performed genotype calling and imputation on this set of target SNPs using read data in BAM format from all 1239 samples using STITCH v1.6.6 (Davies et al., 2016). We ran STITCH in 1 Mb windows in pseudohaploid mode with the following parameters: K=100, nGen=1000, shuffle_bin_radius=100, niterations=40, switchModelIterations=25, buffer=50000. Based on the significant negative relationship between chromosome size and per bp recombination rate in birds (Singhal et al., 2015), STITCH was run across three chromosome classes using the following recombination rate tuning (i.e., >25 Mb, expRate = 1.0; <25 Mb, expRate = 5.0, and < 10 Mb, expRate = 10.0). A final set of 33.16 million SNPs with high information content (INFO_SCORE > 0.4) were retained for downstream analyses (88.2% of the initial set of variants identified). While all 1239 samples were used for improving imputation performance, we only retained a subset of 200 samples (*atropygialis*, N = 48; *cincta*, N = 42, *hecki*, N = 110) in this WGS dataset for downstream analyses (Table S1).

Genotyping the set of 305 samples with ddRAD data was performed in accordance with GATK v4.4 Best Practices (Van der Auwera and O’Connor 2020). An initial set of variants were called for each sample for each chromosome using HaplotypeCaller in GVCF mode, GVCF output from all 305 samples were combined by chromosome using CombineGVCFs, and joint genotyping was performed using GenotypeGVCFs. As above, we restricted downstream analyses to the 30 largest chromosomes in the zebra finch reference. We selected all SNPs on each chromosome with the SelectVariants module of GATK, masked sites that overlapped with annotated repetitive regions, and kept variants that remained after applying the following quality filters: QD < 2, FS > 60.0, MQ < 30.0, ReadPosRankSum < -8.0. We filtered this initial dataset of samples missing data at more than 30% of sites and then of SNPs without data in more than 20% of remaining samples using vcftools v0.1.16 (Danecek et al., 2011). We estimated kinship coefficients between all pairs of individuals using KING v2.3.1 (Manichaikul et al., 2010) and removed two individuals from pairs identified as first-degree relatives. A total of 256 samples (*cincta*, N = 136; *atropygialis*, N = 107; *hecki*, N = 13) and 258,282 SNPs remained in the ddRAD dataset after applying these filtering criteria (Table S3).

### Population structure, genetic differentiation, and summary statistics

We examined population structure and admixture among the samples in our two datasets (i.e., WGS and ddRAD) by performing principal component analysis (PCA), constructing admixture plots, inferring effective migration surfaces, and building maximum likelihood trees. We evaluated the genomic landscape of differentiation among populations by calculating F_ST_ between the three sampled taxa. Each PCA was performed using plink v1.09 (Purcell et al., 2007). For the WGS dataset, we removed SNPs <1 kb apart, with a minor allele frequency less than 0.05, and genotyped in less than 0.95 of samples to retain 854,571 SNPs. For the ddRAD dataset, we removed singleton sites and SNPs <1 kb apart to retain 153,540 SNPs. We then performed PCA sequentially using subsets of the entire dataset. First, using both black-throated finch and long-tailed finch samples. Second, only using samples from the two black-throated finch subspecies. Third, only using samples from the endangered black-throated finch subspecies *cincta*.

We examined geographic structure and admixture patterns among black-throated finch samples using the programs fastStructure v1.0.5 (Raj et al., 2014) and FEEMS v1.0 (Marcus et al., 2021). We restricted these analyses to the ddRAD dataset due to its broader geographic breadth of sampling for black-throated finches than the WGS dataset. Input data for fastStructure was prepared by removing SNPs found in linkage disequilibrium (i.e., r^2^ > 0.2) in 100 kb windows and 10 kb steps using the --indep-pairwise function of plink. A total of 0.2 million SNPs remained after LD pruning. We ran fastStructure with ten cross-validation tests and values of K ranging from 2 to 4. The optimal model complexity was determined using the chooseK.py function of fastStructure. The program FEEMS (Fast Estimation of Effective Migration Surfaces) uses a Gaussian Markov Random Field model to infer and visualize spatially heterogeneous isolation-by-distance patterns (i.e., effective migration rates) on a geographic surface (Marcus et al., 2021). For FEEMS, we generated a genetic distance matrix using the set of LD-pruned SNPs, a geographic distance matrix using the longitude and latitude coordinates of each sample, and defined an outer boundary polygon using https://www.keene.edu/campus/maps/tool/.

We evaluated the genomic landscape of differentiation between *cincta*, *atropygialis*, and *hecki* by calculating F_ST_ between each taxon pair using vcftools v0.1.16. We restricted these analyses to the WGS dataset because it captures a far more complete representation of genomic variation than the ddRAD dataset. We examined F_ST_ in 20 kb sliding windows with 10 kb steps and calculated F_ST_ per SNP after removing singleton sites. For calculations on the Z chromosome, we restricted analysis to males to circumvent any potential issues associated with female hemizygosity. We investigated subspecies specific differentiation patterns within the black-throated finch using the normalized population branch statistic (PBSn1, Malaspinas et al., 2016) in 20 kb sliding windows with the long-tailed finch as outgroup. This modified version of the original PBS statistic rescales by total tree length and has been shown to have a lower false positive rate in identifying local selective sweeps (Shpak et al., 2024).

We assessed the number of private alleles in each taxon and sampling location using a custom perl script from Shogren et al., 2024 (https://github.com/ehshogren/MyzomelaPopulationGenomics). Private alleles were called as monomorphic or biallelic sites observed only in the focal taxon (i.e., *atropygialis*, *cincta*, or *hecki*) and present in at least five individuals. We subsequently scored each location where individuals carrying that private allele were sampled. To quantify the proportion of genetic diversity unique to each sampling locality, we then calculated the proportion of private alleles found in each taxon that were also unique to each sampled population.

### Heterozygosity, relatedness, and inbreeding

We compared genetic diversity in each of our three taxa based on the observed number of heterozygous genotypes in each sample. For the WGS dataset, we estimated mean site-based heterozygosity as the number of heterozygous genotypes on the 29 largest autosomal chromosomes divided by the summed non-repeat masked length of these chromosomes. For the ddRAD dataset, we estimated mean site-based heterozygosity as the total number of heterozygous autosomal genotypes divided by the total number of autosomal SNPs. To account for variation in missingness per sample in the ddRAD dataset, we removed 5 samples missing data at more than 20% of sites and then restricted analyses to the 53,049 SNPs without any missing data across the remaining 251 samples. Counts of heterozygous genotypes were performed using bcftools v1.9 without a minor allele frequency filter. We tested for significant differences in heterozygosity between taxa and between populations within taxa using Tukey’s HSD tests in R (R Core Team, 2022).

We examined variation in relatedness within and between our sampled populations by calculating the pairwise kinship coefficient between all samples in the ddRAD dataset using KING v2.3.1 (Manichaikul et al., 2010). We defined 1^st^ degree (kinship ≥0.18-0.35), 2^nd^ degree (kinship ≥0.09-0.18), and 3^rd^ degree (kinship ≥0.04-0.09) relatives based on the expected distribution of pairwise kinship coefficients for these relationships. Two previously identified 1^st^ degree relatives that had been removed in all other analyses were included in relatedness analyses. We evaluated whether the distribution of pairwise relatedness differed between sampled populations using Tukey’s HSD tests. Three of our populations were comprised of samples collected more than six months apart (*atropygialis*: Red Lily Lagoon in November 2008 and June 2009; *cincta*: Ford’s Dam in November/December 2012 and August/September 2013; Clearwater Dam in October 2008 and June 2009). As we did not observe any significant difference in pairwise relatedness between samples collected from the same location at the same date and samples collected at different dates, we considered all samples from the same location regardless of date collected to come from the same population. Finally, we compared whether the proportion of close relatives (i.e., 3^rd^ degree relatives and closer) observed in each population differed using Pearson’s Chi-squared tests. We restricted this analysis to seven populations with at least 10 individuals sampled to preclude any issues that might result from comparing groups with low sample size. All statistical tests were performed in R (R Core Team, 2022).

We evaluated evidence of inbreeding across populations in our sample set by contrasting the proportion of the genome observed in runs of homozygosity (F_ROH_) due to the established relationship between F_ROH_ and homozygous mutation load (Kardos et al., 2018; Nguyen et al., 2022). We additionally evaluated the number and lengths of runs of homozygosity (ROH) across the genome as the length distribution of ROH can be especially informative about the timing of inbreeding events (Franklin 1977). Specifically, longer ROH are expected to result from more recent episodes of inbreeding while shorter ROH are more indicative of historical bottlenecks that have subsequently been broken up by recombination (Pemberton et al., 2012). We classified ROH and estimated F_ROH_ using autosomal SNPs with the --homozyg function of plink v1.9 with default parameter tuning. We did not perform LD pruning or allele frequency filtering of this dataset because of the downward-biased effect of these filters on estimates of F_ROH_ (Meyermans et al., 2020). We restricted ROH analysis to the WGS dataset because it has a marker density two orders of magnitude greater than the ddRAD dataset (WGS: 25.6 SNPs per kb; ddRAD: 0.26 SNPs per kb).

### Demographic inference

We built maximum likelihood trees including all three taxa using both the WGS and ddRAD datasets with RAxML v8.2.4 (Stamatakis 2014). We used vcftools to generate a set of physically thinned SNP variants (WGS: 10 kb thinned; ddRAD: 5 kb thinned) with a minor allele count of at least two. For the WGS dataset, this resulted in 99,716 SNPs, 51,882 of which (those that had a minor allele in homozygosity in at least one individual) could be used for building a maximum likelihood tree. For the ddRAD dataset, this resulted in 10,632 SNPs, 6,650 could be used for maximum likelihood tree building. For both datasets, we implemented the ASC_GTRGAMMA model in combination with the Lewis correction for SNP ascertainment bias and carried out 350 bootstrap replicates. The resulting phylogenetic relationships between *atropygialis*, *cincta*, and *hecki* were used to frame demographic reconstruction.

We applied the Pairwise Sequential Markovian Coalescent (PSMC; (Li and Durbin 2011) model to infer historical changes in effective population sizes (*N_e_*) in the black-throated finch and long-tailed finch. We restricted analysis to samples with sufficient whole genome sequence data available for PSMC inference (i.e., depth of coverage ≥20×): *atropygialis*, N = 1; *cincta*, N = 1; and *hecki* N = 6 (Table S2). Diploid consensus sequences were generated for each sample using the samtools v1.6 mpileup, bcftools v1.9 call, and the ‘vcf2fq’ function from vcfutils.pl. We only considered the 29 largest autosomal chromosomes, filtered sites with a base and mapping quality below 30, a lower depth of coverage threshold of 10, and an upper depth of coverage threshold equal to twice the mean depth of coverage for each sample. Default parameters were used for PSMC module fq2psmcfa, except that we used a quality filter of 30 and a bin size of 50 bp, to account for the higher density of heterozygous sites in finches compared to humans. Following the recommendations of Hilgers et al. (2024), we adjusted default tuning of the number of time intervals to avoid false population size peaks/crashes and ran PSMC with the following parameters: -N30, -t5, -r5, -p “2+2+25*2+4+6”. We converted evolutionary units to millions of years and effective population size (*N_e_*) using a generation time of two years estimated from equation 2 of Bird et al. (2020) based on the age of first reproduction in the wild (1 year) and maximum longevity (8 years). We used a mutation rate (per site, per generation) of 5.85 × 10^-9^ based on the estimated germline mutation rate of the zebra finch (Bergeron et al., 2023). We focused our evaluation on the relative differences in demographic trends between taxa as these (i.e., population bottlenecks and historical decline) are not affected by parameter tuning, which are otherwise extremely sensitive to assumptions about the mutation rate and generation time.

Lastly, we examined the joint demographic history within the black-throated finch with fastsimcoal2 v27 (Excoffier et al., 2013). We performed demographic inference at two timescales. In the first, focused on the dynamics of subspecies, we modeled the demographic histories of *atropygialis* and *cincta*. In the second, focused on the dynamics within subspecies *cincta*, we modeled the demographic histories of remaining population stronghold from the Townsville Coastal Plain and the Galilee Basin. For both analyses, we used our ddRAD dataset, built two-dimension site frequency spectra (2D-SFS) using easySFS (https://github.com/isaacovercast/easySFS), and utilized hypergeometric down projection of the allelic sample size to maximize the number of segregating sites per lineage. The resulting 2D-SFS were based on autosomal variants that had been thinned so that no two sites were within 1 kb: resulting in 18,946 SNPs, *atropygialis* and *cincta*; and 18,943 SNPs, Townsville Coastal Plain and Galilee Basin. We tested an initial suite of two-population models that encompassed 12 scenarios that varied in a) the timing of divergence, b) the number of distinct migration rates, c) the number and nature of population size fluctuations, and d) the size of ancestral populations. We performed 100 independent parameter runs per model using the following options: -n 100000 -L 100 -C 1 -y 5 -0 -m -q –logprecision 18 –brentol 0.0001. As above, we used a generation time of two years and an approximate autosomal germline mutation rate of 5.85 × 10^-9^ (per site, per generation). We compared model fit based on the maximum likelihood estimate distribution of the top five independent runs for each model and then selected the best-fit model using the Akaike Information Criterion (AIC) (Bozdogan 1987). We randomly selected sets of 20,150 SNPs (∼10% of total autosomal SNPs) to generate 100 non-parametric bootstrap 2D-SFS datasets using easySFS. We obtained confidence intervals for parameter estimates under the best-fitting model after running a further 50 replicates per simulated dataset (i.e., for a total of 5000 parameter estimates).

## RESULTS

### Population structure and genetic differentiation

We recovered three clearly distinct population clusters from PCA, which correspond to northern and southern black-throated finch subspecies (*atropygialis* and *cincta*, respectively) and the long-tailed finch (subspecies *hecki*; Figure 1B). Both the WGS and ddRAD datasets recovered the separation of *hecki* from *atropygialis* and *cincta* along PC1 with subsequent separation of *atropygialis* from *cincta* along PC2 (Figure 1B and Figure S1). These three population clusters effectively recapitulate the geographic distribution of the taxa from which individuals were sampled (Figure 1). The clustering of individuals is consistent with two biogeographic barriers to gene flow: the east-to-west Carpentarian Gap (CG) between the long-tailed finch and black-throated finch and the northeast-to-southwest Einasleigh Uplands (EU) between *atropygialis* and *cincta* (Figure 1). Considering black-throated finch samples in isolation, we observed further evidence of a north-south genetic break between *atropygialis* and *cincta* samples along PC1 with no evidence of any recently admixed individuals (Figure 1C and Figure S1). Within *cincta*, PCA suggests a genetic break exists between the Townsville Coastal Plain and the Galilee Basin (Figure 1C), which we discuss further below. The genetic distinctiveness of *atropygialis* and *cincta* was additionally supported in a geographically agnostic assessment of population structure (K = 2) where nearly all samples had admixture coefficients less than 5% (*atropygialis*: 103 of 107 samples; *cincta*: 131 of 136 samples; Figure 2A). The longitude and latitude coordinates of where a sample was collected explained nearly all variation in PC1 loading (r^2^ = 0.94) and admixture proportion (r^2^ = 0.90) as tested by linear regression. Furthermore, we found evidence of a geographic barrier to gene flow between populations of *atropygialis* and *cincta* associated with lower effective migration rates (*w*) across the Einasleigh Uplands (Figure 2B). This upland region, composed primarily of eroded volcanic rock, has previously been described as a biogeographic barrier in some twenty avian species – including the black-throated finch – with range edges, stepped phenotypic clines, or hybrid zones across this region (Keast 1961; Ford 1986).

**FIGURE 2.**
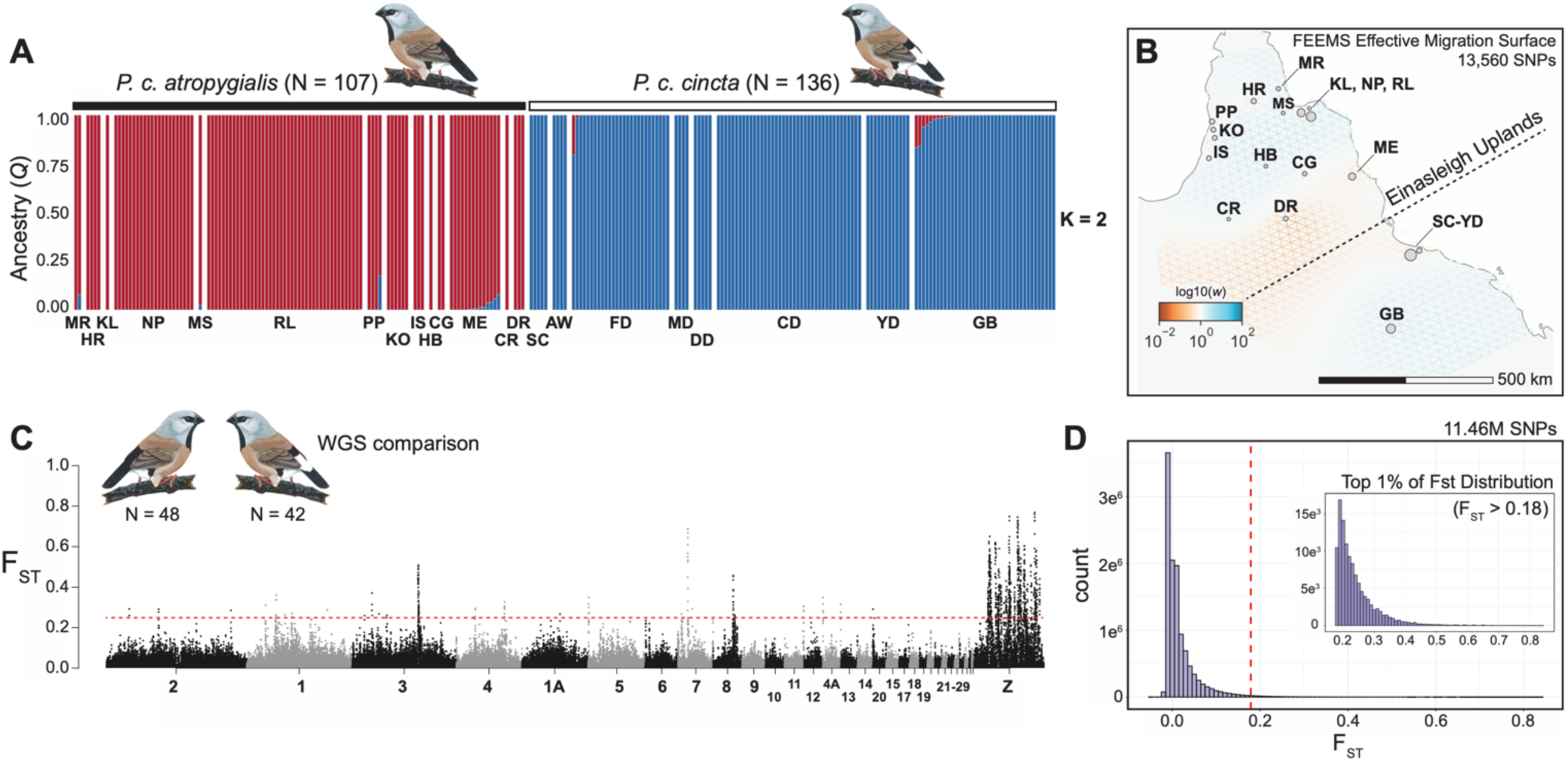
Population structure and genomic differentiation in the black-throated finch. (A) Geographic structure in the black-throated finch is associated first with subspecies identity. The ancestry coefficient (Q) was inferred per sample using fastSTRUCTURE (K = 2) for *atropygialis* (N = 107) and *cincta* (N = 136). (B) The effective migration surface of the black-throated finch was estimated using FEEMS (Marcus et al., 2021). Effective migration rates are color-coded on a log scale from low migration rate (red tones) to high migration rate (blue tones) against a model of isolation by distance. The approximate location of the Einasleigh Uplands is annotated over the inferred historical barrier to migration between subspecies. (C) Genomic differentiation (F_ST_) between *atropygialis* (N = 48) and *cincta* (N = 42) in 20 kb windows with 10 kb steps. The dashed red horizontal line represents the 99^th^ percentile threshold for the window based F_ST_ distribution (i.e., F_ST_ = 0.22). (D) Histogram of F_ST_ per-site between black-throated finch subspecies following singleton site removal. The dashed red vertical line represents the 99^th^ percentile threshold for the site based F_ST_ distribution (i.e., F_ST_ = 0.18). Inset histogram panel depicts the distribution of the top 1% most differentiated SNPs (most differentiated SNP, chr8:21796958, F_ST_ = 0.84). Results in panels (A) and (B) based on the ddRAD dataset and panels (C) and (D) on the WGS dataset.

Despite clear evidence of geographic structure, the magnitude of genomic differentiation between *atropygialis* and *cincta* was low (mean weighted-F_ST_ = 0.029) but was on average 4× greater on the Z chromosome than on the autosomes (mean autosomal F_ST_ = 0.025; mean chrZ F_ST_ = 0.082; Figure 2C). None of the 11.46 million non-singleton SNPs examined were observed as fixed differences between subspecies (Figure 2D). The most differentiated SNPs (84 SNPs with F_ST_ > 0.75) were clustered into six genomic regions of elevated differentiation (Table 1). Only one differentiation peak was located on an autosome (chromosome 8) while the other five were located on the Z chromosome. We identified 6 protein-coding genes within 20 kb of these differentiation peaks (median = 1; range = 0 to 2 genes; 76% of highly differentiated SNPs were intergenic). In total, 15% of highly differentiated SNPs were found in putatively *cis*-regulatory regions (i.e., within 20 kb of genes), 4% in intronic regions, and 5% in coding regions. Three of the six differentiation peaks between black-throated finch subspecies could be attributed to large allele frequency changes since their divergence in *cincta*, two to *atropygialis*, and one to both subspecies (Table 1). In *atropygialis*, the strongest signature of positive selection was on chromosome 8 and associated with the tyrosine phosphatase receptor type C gene *PTPRC. PTPRC* – also known as *CD45* – is a large transmembrane glycoprotein found on the cell surface of all hematopoietic cells other than mature erythrocytes and is involved in innate immune response (Al Barashdi et al., 2021). Recent work in the long-tailed finch reported evidence of a selective sweep on *PTPRC* in subspecies *acuticauda* (Hooper et al., 2024). In *cincta*, the strongest signature of selection was located on the Z chromosome in a region that encompassed the polyadenylate-binding protein-interacting protein 1 gene *PAIP1* and the mitonuclear associated NAD(P) transhydrogenase gene NNT.

**Table 1.**
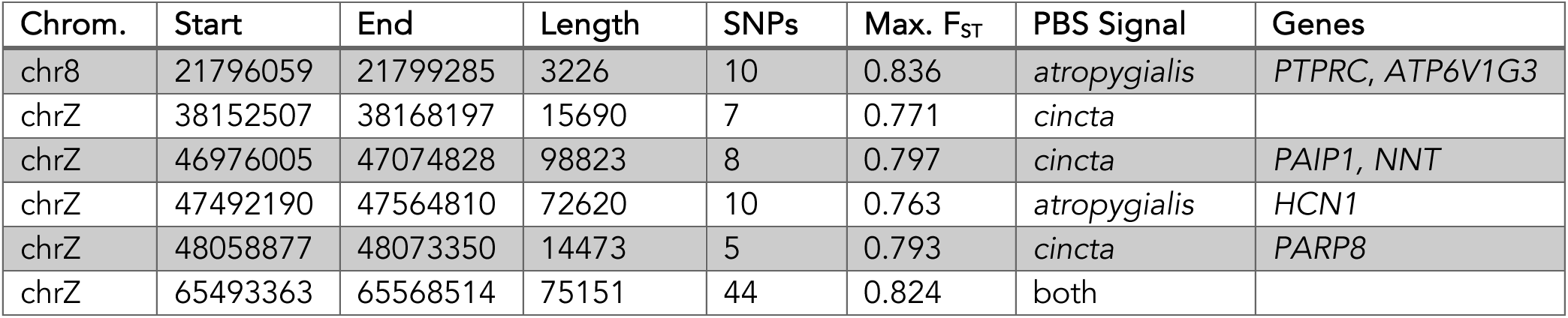
Summary of differentiation peak regions observed between black-throated finch subspecies. We polarized differentiation to the subspecies of origin using the normalized population branch statistic (PBSn1). Genes within 20 kb of the start and end coordinates of each peak region are listed. See full PBS results in Table S5.

| Chrom. | Start | End | Length | SNPs | Max. F<sub>ST</sub> | PBS Signal | Genes |
| --- | --- | --- | --- | --- | --- | --- | --- |
| chr8 | 21796059 | 21799285 | 3226 | 10 | 0.836 | *atropygialis* | *PTPRC*, *ATP6V1G3* |
| chrZ | 38152507 | 38168197 | 15690 | 7 | 0.771 | *cincta* |  |
| chrZ | 46976005 | 47074828 | 98823 | 8 | 0.797 | *cincta* | *PAIP1*, *NNT* |
| chrZ | 47492190 | 47564810 | 72620 | 10 | 0.763 | *atropygialis* | *HCN1* |
| chrZ | 48058877 | 48073350 | 14473 | 5 | 0.793 | *cincta* | *PARP8* |
| chrZ | 65493363 | 65568514 | 75151 | 44 | 0.824 | both |  |

Genomic differentiation between the long-tailed finch and each black-throated finch subspecies was high (mean F_ST_ *atr-hec* = 0.226; *cin-hec* = 0.243) and was on average ∼3× greater on the Z chromosome (mean autosomal F_ST_ *atr-hec* = 0.192 and *cin-hec* = 0.211; mean chrZ F_ST_ *atr-hec* = 0.641 and *cin-hec* = 0.645). We observed 233,260 highly differentiated SNPs (F_ST_ > 0.75) between the long-tailed finch and the black-throated finch – both *atropygialis* and *cincta* – and 73,863 of these were observed as fixed differences (62.2% of which were Z-linked). Genetic differentiation is elevated across the entire genome (Figure S2), suggesting a substantial role of allopatric divergence between these two species.

### Fine-scale genetic structure within the southern black-throated finch

We found evidence of strong spatial genetic structure between populations of subspecies *cincta* collected from the Townsville Coastal Plain. Principal component analysis recovers a characteristic isolation-by-distance signature between populations separated by two landscape features: the Townsville suburb of Kelso and the Ross River Dam (Figure 3). Akin to the genetic structure expected of a ring species (Irwin et al., 2001), genetic distance was low between population pairs moving clockwise around the Ross River Dam until comparing populations on opposite sides of Kelso (Figure 3D). Indeed, the greatest genetic distance of populations sampled in the Townsville region was observed between individuals from Sunbird Creek (SC) and those from Clearwater Dam (CD) (Figure 3D). Although these populations are located only 8.6 km apart, they are separated by both a suburb and a water body, two features that appear to constitute major barriers to dispersal (Figure 3A). Remarkably, all Townsville populations carried alleles private to that sampling locality: on average, 5.7% of autosomal variants and 6.6% of Z-linked SNPs segregating only in *cincta* were unique to each population; Table S6). All individuals from the Galilee Basin, located ∼275 km southwest of Townsville and constituting the only other major population of *cincta*, were recovered as a distinct genetic cluster (Figure 3D). Moreover, 32.5% of autosomal and 35.1% of Z-linked SNPs only found in *cincta* were private to the Galilee Basin, highlighting the genetic distinctiveness of this geographic region (Table S6).

**FIGURE 3.**
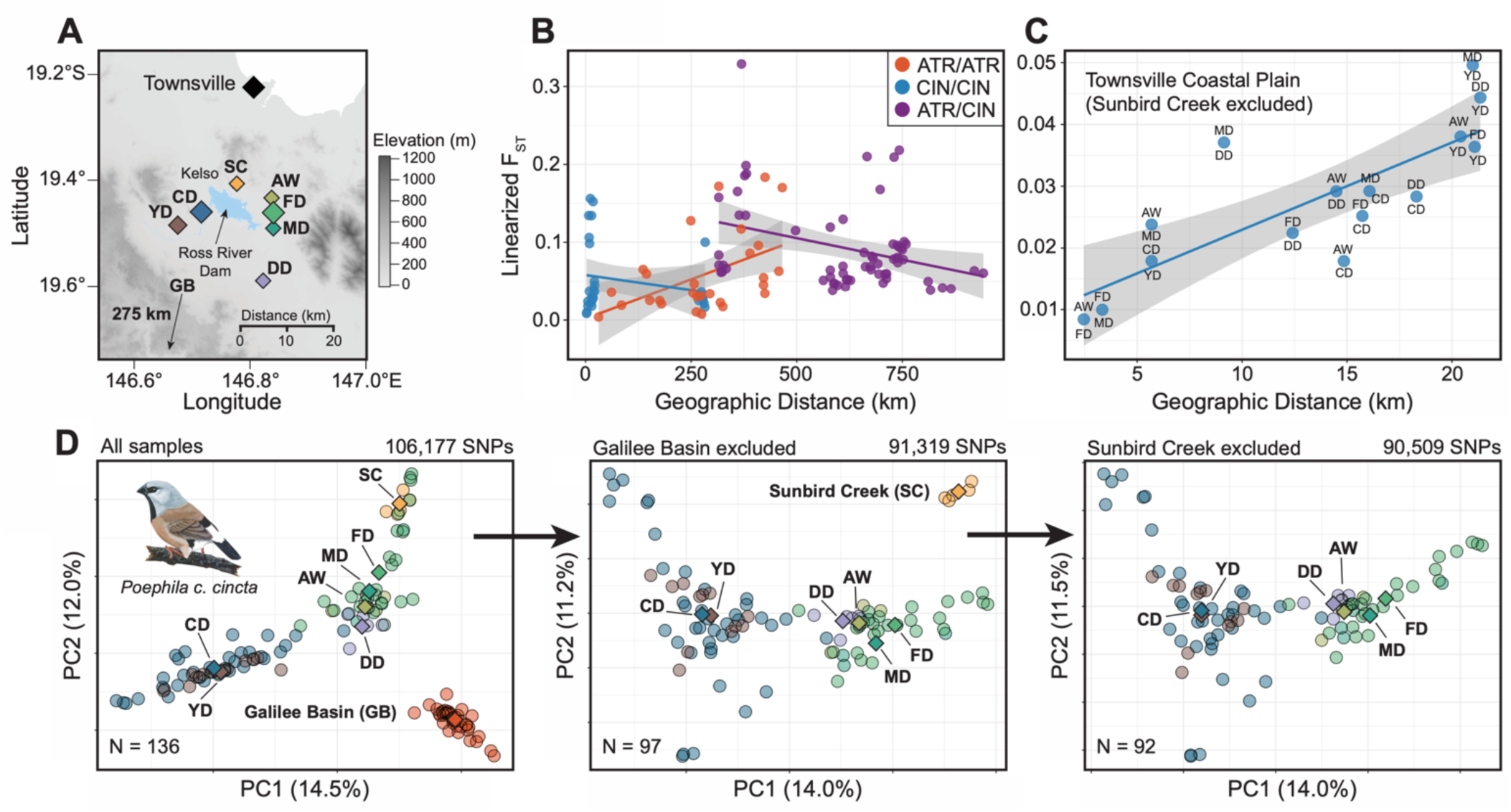
Fine-scale genetic structure within black-throated finch (subspecies *cincta*) populations of the Townsville coastal plain, QLD. (A) Geographic distribution of seven sampled populations, Ross River Dam, suburb of Kelso, and Townsville city center are labeled. Topographical relief is represented in greyscale with higher elevations in darker colors. For reference, the distance and direction to a population of *cincta* located in the Galilee Basin (GB) is indicated on the map. (B) Isolation by distance (IBD) between populations within lineages (*atropygialis* in red; *cincta* in blue) and between lineages in purple. The linear relationship between geographic distance and genetic distance is shown for each group with standard error represented in grey. (C) IBD between *cincta* populations of the Townsville coastal plain. The pair of populations compared is indicated for each point. (D) Principal components analysis of 136 *cincta* samples reveals evidence of fine-scale geographic structure between populations of the Townsville coastal plain and of the Galilee Basin (GB) located 275 km to the southwest. Population centroids, representing the mean loading of PC1 and PC2, are plotted and labeled for reference. The most divergent population was sequentially excluded from left to right panels.

### Heterozygosity, relatedness, and inbreeding

We observed significant differences in individual heterozygosity between black-throated finch subspecies and the long-tailed finch (Figure 4; Table 2). Mean site-based heterozygosity was lowest in *cincta* (mean and standard deviation, WGS: 0.0037 ± 0.0001; ddRAD: 0.0406 ± 0.0046), slightly but significantly greater in *atropygialis* (WGS: 0.0038 ± 0.0001; ddRAD: 0.0430 ± 0.0046), and greatest in *hecki* (WGS: 0.0045 ± 0.0001; ddRAD: 0.0507 ± 0.0020) (Table 2; Figure 4). Individual estimates of heterozygosity were strongly correlated between the WGS and ddRAD datasets (r^2^ = 0.65, N = 82) as tested by linear regression. The reported difference in heterozygosity between *atropygialis* and *cincta* appears to be driven by two Townsville area populations west of the Ross River Dam (CD and YD; Figure 4D). There was no significant difference in heterozygosity between *atropygialis* and *cincta* when these two populations were removed.

**FIGURE 4.**
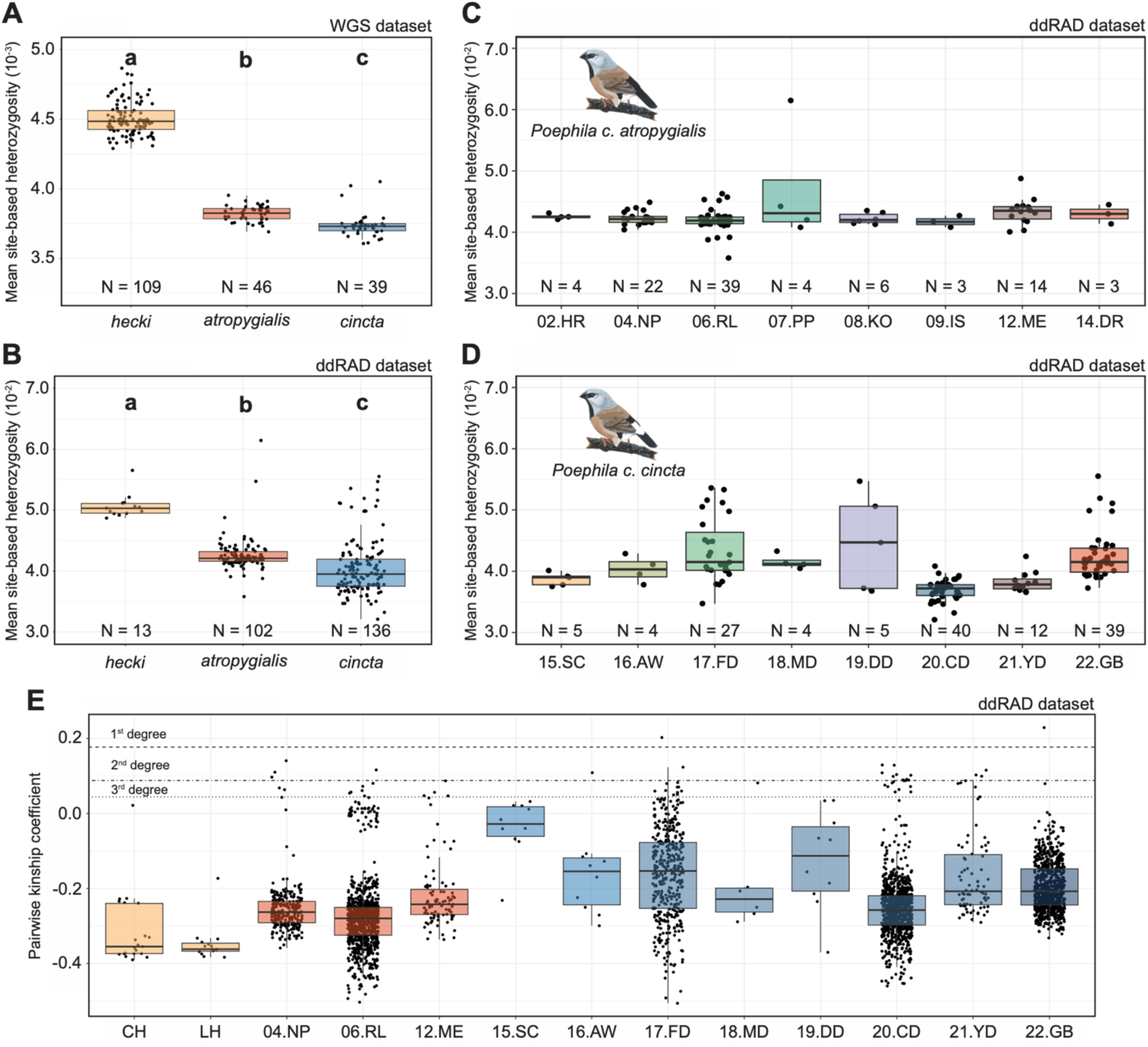
Variation in heterozygosity and relatedness. (A) Mean site-based observed heterozygosity in *hecki*, *atropygialis*, and *cincta* based on WGS dataset and (B) ddRAD dataset. The bottom, bolded middle, and topmost lines of the boxplots represent the first quartile, mean, and third quartile, respectively. Each point represents an individual. (C) Mean site-based observed heterozygosity in *atropygialis* and (D) in *cincta* by population. Only populations with at least 4 individuals are shown. (E) Mean pairwise relatedness (kinship coefficient) between individuals in populations from *hecki* (yellow), *atropygialis* (red), and *cincta* (blue). Each point represents a pair of individuals. Dashed lines represent the kinship threshold for first-, second-, and third-degree relatives. Different letters in (A) and (B) indicate statistical significance between taxa (adjusted P < 0.05; Tukey’s HSD test).

**Table 2.** Mean site-based observed heterozygosity by taxon and by population. Population ‘All’ represents the mean for all samples in each taxon. Only results from populations with at least six samples are shown; see full results in Tables S6-S9.

| Taxon | Population | Longitude | Latitude | ddRAD |  | WGS |  |
| --- | --- | --- | --- | --- | --- | --- | --- |
|  |  |  |  | N | Heterozygosity | N | Heterozygosity |
| *atropygialis* | All | - | - | 102 | 0.0430 | 46 | 0.00382 |
|  | Nifold Plains (NP) | 143.97 | -14.64 | 22 | 0.0423 | 16 | 0.00384 |
|  | Red Lily Lagoon (RL) | 144.16 | -14.85 | 39 | 0.0429 | 16 | 0.00381 |
|  | Kowanyama (KO) | 141.70 | -15.36 | 6 | 0.0423 | - | - |
|  | Mareeba (ME) | 145.35 | -16.93 | 14 | 0.0434 | - | - |
| *cincta* | All | - | - | 136 | 0.0406 | 39 | 0.00373 |
|  | Ford's Dam (FD) | 146.86 | -19.50 | 27 | 0.0435 | - | - |
|  | Clearwater Dam (CD) | 146.71 | -19.46 | 40 | 0.0370 | 21 | 0.00373 |
|  | Yellow Dam (YD) | 146.66 | -19.48 | 12 | 0.0382 | 18 | 0.00372 |
|  | Galilee Basin (GB) | 146.26 | -21.92 | 39 | 0.0426 | - | - |
| *hecki* | All | - | - | 13 | 0.0507 | 109 | 0.00451 |
|  | Maryfield | 133.40 | -15.83 | - | - | 15 | 0.00450 |
|  | October Creek | 134.86 | -16.63 | - | - | 41 | 0.00456 |
|  | MacArthur River | 135.87 | -16.51 | - | - | 13 | 0.00442 |
|  | Chidna | 139.45 | -19.45 | 7 | 0.0509 | 20 | 0.00447 |
|  | Leichardt | 139.74 | -20.26 | 6 | 0.0504 | 14 | 0.00449 |

We found clear differences in pairwise relatedness between individuals from populations of *atropygialis*, *cincta*, and *hecki* (Figure 4E). Consistent with expectations for larger and more stable populations, pairwise relatedness tended to be lower between individuals from populations of *atropygialis* (mean and standard deviation, kinship coefficient -0.268 ± 0.089) than between individuals from populations of *cincta* (-0.210 ± 0.095). Mean pairwise relatedness was significantly lower in *hecki* (-0.328 ± 0.081) than in both *cincta* and *atropygialis* (adjusted *P* < 0.005 for both contrasts, Tukey’s HSD test). Consistent with the fine-scale genetic structure described above, mean pairwise relatedness differed substantially between populations of *cincta* sampled around the Townsville Coastal Plain (Figure 4E), suggesting limited dispersal between locations, sampling stochasticity, or both processes. We did not, however, observe a significant difference in the overall proportion of close relationships (i.e., pairs with kinship coefficients ≥0.044 indicating 3^rd^ degree relatives – first cousins – or greater) between *atropygialis* and *cincta* (Pearson’s Chi-squared test: *X*^2^ = 1.66, *df* = 1, *p* = 0.1975). Of the 45 pairs of close relatives identified from the Townsville Coastal Plain, only three pairs represented individuals collected from different sampling locations (Table 3 and Table S11). All three pairs were 3^rd^-degree relative relationships that included individuals collected from Ford’s Dam (FD) and Dotterel Dam (DD), two populations separated by 8.1 km.

**Table 3.** Proportion of closest pairwise relationships (those with kinship coefficients ≥ 0.044 indicating 3^rd^ degree relatives or greater) within each population. Thresholds for each relationship class were based on established expectations for the distribution of kinship coefficients (Manichaikul et al., 2010). The number and proportion of pairwise relationships belonging to each class are reported. See full results in Table S10.

| Taxon | Population | N | Close relatives | 3<sup>rd</sup> degree | 2<sup>nd</sup> degree | 1<sup>st</sup> degree |
| --- | --- | --- | --- | --- | --- | --- |
| *atropygialis* | Nifold Plains (NP) | 23 | 5 (2.0%) | 2 (0.8%) | 3 (1.2%) | - |
|  | Red Lily Lagoon (RL) | 43 | 7 (0.8%) | 6 (0.7%) | 1 (0.1%) | - |
|  | Mareeba (ME) | 14 | 4 (4.4%) | 4 (4.4%) | - | - |
| *cincta* | Sunbird Creek (SC) | 5 | 0 (0.0%) | - | - | - |
|  | Anthill West (AW) | 5 | 1 (10.0%) | - | 1 (10.0%) | - |
|  | Ford's Dam (FD) | 29 | 12 (3.0%) | 10 (2.5%) | 1 (0.2%) | 1 (0.2%) |
|  | Mango Dam (MD) | 4 | 1 (16.7%) | 1 (16.7%) | - | - |
|  | Dotterel Dam (DD) | 5 | 0 (0.0%) | - | - | - |
|  | Clearwater Dam (CD) | 43 | 20 (2.2%) | 10 (1.1%) | 10 (1.1%) | - |
|  | Yellow Dam (YD) | 13 | 8 (10.3%) | 5 (6.4%) | 3 (3.8%) | - |
|  | Galilee Basin (GB) | 40 | 2 (0.3%) | 1 (0.1%) | - | 1 (0.1%) |
| *hecki* | Chidna | 7 | 0 (0.0%) | - | - | - |
|  | Leichardt | 6 | 0 (0.0%) | - | - | - |

We detected 12 ROH on five of the twenty-nine autosomal chromosomes examined (Table S12). In contrast to expectations based on differences between taxa in heterozygosity and relatedness, all ROH were identified in *hecki* and ranged in length from 1.0 to 1.9 Mb. Eleven individuals had a single ROH, and one individual had two. Nine of the identified ROH shared overlapping coordinates across the individuals in which they were observed, a signature more suggestive of background selection than inbreeding (Table S12).

### Demographic inference

Phylogenetic analyses supported a deep split between a clade consisting of all black-throated finch individuals and a clade with all long-tailed finch individuals (Figure 5A and 5B). All *cincta* individuals formed a monophyletic clade with full bootstrap support (i.e., 1.0) in the WGS tree, while all but one *cincta* individual formed a monophyletic clade with low bootstrap support (i.e., <0.5) in the ddRAD tree. Both datasets recovered *atropygialis* as a paraphyletic clade containing the *cincta* clade (Figure 5A and 5B).

**FIGURE 5.**
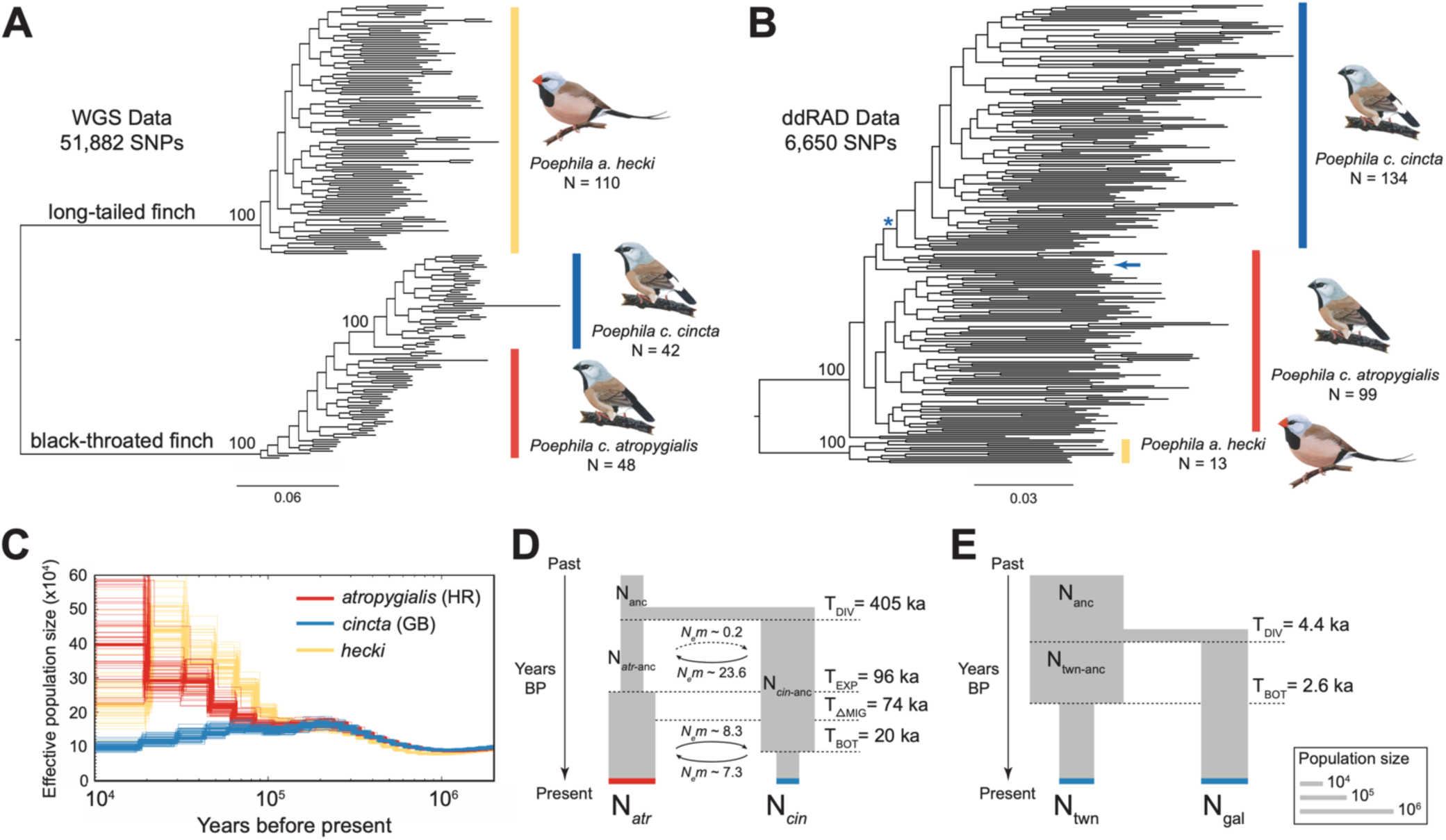
Phylogenetic relationships and demographic reconstruction of the black-throated finch and long-tailed finch. Maximum-likelihood phylogenetic trees constructed with RAxML based on (A) 51,882 SNPs in the WGS dataset and (B) 6,650 SNPs in the ddRAD dataset. Only nodes with 100% bootstrap support are labeled. Vertical colored bars denote the tips belonging to each taxon: *hecki* (yellow), *cincta* (blue), and *atropygialis* (red). In (B) the blue asterisk denotes the clade comprising all *cincta* samples except for the one sample, noted with a blue arrow. (C) Effective population sizes (N_e_) inferred from representative *hecki* (yellow), *atropygialis* (red), and *cincta* (blue) genomes using PSMC, assuming a generation time (*g*) of 2 years and a germline mutation rate (μ) of 5.85e^-09^ per site per generation. Representative samples came from: Holroyd River (HR), *atropygialis*; Galilee Basin (GB), *cincta*; and October Creek, *hecki*. (D) Best-fit demographic model of black-throated finch subspecies *atropygialis* and *cincta* and (E) of *cincta* populations from the Townsville Coastal Plain (twn) and Galilee Basin (gal). Estimates of effective population size (*N_e_*), migration (*m*), and the timing of divergence (T_DIV_), bottleneck (T_BOT_), and expansion (T_EXP_) events are labeled in thousands of years (ka).

Demographic inference based on pairwise sequential Markovian coalescent (PSMC) analyses suggests that black-throated finch lineages ancestral to subspecies *atropygialis* and *cincta* began diverging from each other approximately 100 kya (Figure 5C). Following this divergence event, however, each black-throated finch subspecies exhibits highly discordant demographic histories: the inferred effective population size (*N_e_*) of *cincta* has been low and gradually in decline, while *atropygialis* exhibits evidence of sustained increase towards the present day (Figure 5C). The long-tailed finch and black-throated finch appear to have diverged over 1 mya with the two taxa sharing similar demographic trajectories until around 100 kya, with *hecki* showing evidence of an increase in *N_e_* until approximately 20 kya (Figure 5C).

Lastly, we used the joint site frequency spectrum to infer best-fit demographic models to describe the evolutionary histories of 1) black-throated finch subspecies *atropygialis* and *cincta* and 2) the two remaining population centers of *cincta*: Townsville Coastal Plain and the Galilee Basin. The black-throated finch subspecies data were best fit by a model with a long history of high migration: initial migration rates of 1.8 × 10^-4^ (*atr*-to-*cin*, 95% confidence interval (CI) = 1.3 × 10^-9^ to 5.6 × 10^-3^) and 4.4 × 10^-5^ (*cin*-to-*atr*, 95% CI = 1.3 × 10^-9^ to 4.9 × 10^-3^) after a population split 0.41 (95% CI = 0.02 to 1.91) million years ago (Figure 5D; Table S13). As the estimated effective population sizes (*N_e_*) are large (*atropygialis N_e_* = 2.6 × 10^4^ to 1.1 × 10^6^; *cincta N_e_* = 5.1 × 10^3^ to 2.8 × 10^5^), the effective number of migrants per generation (*N_e_m*) has been consistently high over time: *N_e_m* = 8.3 (*atr*-to-*cin*) and *N_e_m* = 7.3 (*cin*-to-*atr*), with a shift to a highly asymmetrical level of migration >74,000 years ago when *N_e_m* < 1 (*atr*-to-*cin*) and *N_e_m* > 20 (*cin*-to-*atr*) (Figure 5D). A strongly asymmetrical rate of migration, consistent with the results of phylogenetic analyses finding that *cincta* is a nested subclade within *atropygialis*, may be due to the maintenance of ancestral polymorphisms in the larger population (Figure 5A and 5B). Consistent with PSMC results, the best fit model suggests that both subspecies have experienced population size fluctuations: expansion in *atropygialis* starting ∼96,000 years ago and contraction in *cincta* in the last 20,000 years (Figure 5D). The two remaining population centers of the southern black-throated finch were best fit by a demographic model with no migration and variable population size in the Townsville Coastal Plain population (Figure 5E; Table S14). After an estimated split between these two populations ∼4,400 (95% CI = 1,868 to 6,116) years ago, Townsville Coastal Plain *cincta* have experienced a population decline over the last ∼2,550 (95% CI = 804 to 5,394) years. The estimated effective population size of Townsville Coastal Plain *cincta* was inferred to be approximately 50% of the size of Galilee Basin *cincta* (twn: N_e_ = 26,802 to 162,333; gal: N_e_ = 73,467 to 245,596) (Figure 5E).

## DISCUSSION

Habitat fragmentation and degradation due to accelerating anthropogenic modification of the environment pose significant threats to biodiversity around the world (Brooks et al., 2002; Hanski 2011; Diaz et al., 2019). Species restricted to tropical grassland and savanna habitats are particularly at risk from land use change for agriculture, grazing, urbanization, and mining (Franklin 1999; Laurance et al., 2011; Parr et al., 2014; Garnett and Baker, 2021). Effective conservation management plans require thorough evaluations of genetic diversity and demographic history to safeguard the adaptive resilience of species to anthropogenic habitat loss and climate change (Exposito-Alonso et al., 2022). In this study, we report on the geographic structure, genetic distinctiveness, and demographic history of an endangered songbird native to the tropical savanna grasslands of Queensland, Australia: the black-throated finch. We find that northern and southern subspecies are genetically distinct, geographically isolated by a natural biogeographic barrier to gene flow known as the Einasleigh Uplands, and despite signatures of gene flow following divergence they last shared a common ancestor over 100,000 years ago. Within the southern black-throated finch, we find clear evidence of genetic differentiation between samples from the Galilee Basin and the Townsville Coastal Plain, the two remaining population centers of this endangered finch. Moreover, we report microgeographic genetic structure from the Townsville Coastal Plain between populations <20 km apart that appears to be associated with barriers to dispersal caused by anthropogenic habitat modification over the last 50 years. We elaborate on each of these findings and how the demographic history and geographic structure of the black-throated finch informs conservation management priorities for this species below.

While the biogeography of speciation of the Australian avifauna has historically been better studied than in any other continental assemblage of birds (Ford 1974; Keast 1981; Cracraft 1986), species limits and the dynamics of gene flow between many such lineages are far from settled (Peñalba et al., 2019; Joseph 2021). Finches in the genus *Poephila*, subject of numerous biogeographic studies and one of the earliest molecular efforts to investigate the demography and speciation history of a vertebrate species assemblage (Jennings and Edwards 2005), are no exception (Hooper et al. 2019; Lopez et al., 2021). In this study, we focused first on the significance of two biogeographic barriers in the divergence histories of *Poephila* sister species: the black-throated finch and long-tailed finch. Both the Carpentarian Gap, at the species level, and the Einasleigh Uplands, at the subspecies level, appear to have been historically effective barriers to gene flow between *Poephila* lineages (Figure 1). The Carpentarian Gap, an arid zone barrier between the open savanna habitat favored by many species of Australian birds (Keast 1981), appears to have been an effective barrier to migration between long-tailed finches and black-throated finches. A model of allopatric speciation best fits our lack of evidence of admixture across this biogeographic barrier (Figure 1), an elevated F_ST_ landscape genome-wide (Figure S2), the recovery of a deep phylogenetic split between the two species (Figure 5), and a 1.75 (95% HPD interval: 1.28 to 2.28) million years ago divergence estimate based on mitochondrial data (Lopez et al., 2021).

In contrast, the Einasleigh Uplands, a region of volcanic uplift in northern Queensland, appears to have functioned as a somewhat porous barrier to gene flow between black-throated finch subspecies. Demographic analyses based on the pairwise sequential Markovian coalescent and site frequency spectrum indicate that the northern and southern black-throated finch last shared a common ancestor 0.1-0.4 million years ago, respectively, but diverged with gene flow for most of their subsequent history (Figure 5). While we did not observe any evidence of recent admixture between *atropygialis* and *cincta* in our sample set (Figure 1 and Figure 2), and the extant contracted range of *cincta* would appear to preclude this possibility today, the two subspecies were historically reported to hybridize where their ranges came into contact within this century (Keast 1958). We found that each subspecies harbors many private alleles (*atropygialis*: 25.9% of variants; *cincta*: 17.4% of variants), but we observed no fixed SNP differences between them – likely due to a combination of a large ancestral effective population size and evidence of their divergence with gene flow. Despite the difference in rump color between *cincta* (white rump, ancestral state) and *atropygialis* (black rump, derived state), the key phenotypic difference between these two taxa, none of the genes located within the top 99^th^ percentile of the population branch statistic distribution for *atropygialis* had an affiliation with the melanogenesis pathway. As such, the genetic basis of this key phenotypic difference remains to be identified. In contrast to their low level of genetic differentiation, the long-term demographic trends of each subspecies are quite distinct: population expansion in *atropygialis* and population decline in *cincta*. Notably, signatures of population decline in the southern black-throated finch precedes the anthropogenic habitat change over the last two centuries following European arrival and suggests that this subspecies may have been more susceptible to the effects of these changes, as we discuss further below.

We seek to highlight two important findings regarding geographic structure and genetic diversity within the southern black-throated finch. First, we find that the two remaining population centers of this endangered finch, located in the Galilee Basin and on the Townsville Coastal Plain, are genetically distinct from each other (Figure 3). Each population harbors substantial genetic variation unique to subspecies *cincta* that is not present in the other: Galilee Basin: 32.5% of private alleles; Townsville: 51.4% of private alleles. Demographic modeling suggests that meaningful levels of gene flow between populations ceased ∼4,000 years ago and that Townsville Coastal Plain *cincta* have experience a population decline over the last ∼2,500 years (Figure 5E). Owing largely due to its proximity to human occupation, the ecology and demography of the Townsville Coastal Plain population has been studied to a far greater degree than the Galilee Basin population (Mula Laguna et al., 2019). The identified genetic distinctiveness of these two meta-populations, combined with the unique combination of anthropogenic challenges they face (e.g., mining versus urbanization), complicates the long-term preservation of the southern black-throated finch and necessitates further ecological and demographic research in both locations.

Second, we find evidence of microgeographic genetic structure between southern black-throated finches from the Townsville Coastal Plain (Figure 3). Remarkably, genetic structure appears to be associated in a ring-species like arrangement around the reservoir of the Ross River Dam: genetic distance increases linearly with geographic distance around the reservoir with a major break in the northwest between populations on opposite sides of the Townsville suburb of Kelso (pops. SC and CD, Figure 3). The Ross River Dam was constructed in 1971 to manage annual flooding of the Ross River and likely constitutes a major water barrier to dispersal in these finches. Evidence of fine-scale genetic structure between Townsville Coastal Plain populations, first reported in a dissertation chapter by Tang (2016) based on a suite of 13 microsatellite markers, was attributed to extensive habitat fragmentation in this region. We examined the same set of samples as this prior study, now represented with genomic data, and further strengthen support for this microgeographic genetic structure. Monitoring efforts from the Townsville Coastal Plain indicate that the southern black-throated finch is generally rather sedentary, with mean home ranges varying between 12 hectares during the breeding season and 51 hectares during the non-breeding season (Rechetelo 2016). Long-distance dispersal appears to be rare, most individuals remained within 200 m of where they were originally banded, although a few individuals were observed to have dispersed more than 15 km in less than 50 days (Rechetelo 2016). Consistent with results from monitoring efforts, we did not observe any first- or second-degree relatives between sampled populations, although the three third-degree relative pairs that we did detect (between pops. FD and DD, separated by 8.1 km) indicate that immigration is a non-negligible factor in the demography of the Townsville Coastal Plain.

Despite the striking microgeographic genetic structure observed in southern black-throated finches from the Townsville Coastal Plain, we did not detect evidence of any major genetic consequences of population bottlenecks (e.g., elevated levels of inbreeding or an enrichment of runs of homozygosity). However, variation in observed heterozygosity and pairwise relatedness was greater in these populations than observed in *atropygialis* or *cincta* from the Galilee Basin (Figure 4). This suggests that these small and isolated populations are already highly susceptible to genetic drift and any stochastic processes that might further reduce genetic variation or persistence. Immigration, for example, plays a key role in the dynamics of local bird populations (Chen et al., 2016; Millon et al., 2019; Summers et al., 2024) and the cessation of immigration can result in local extirpation even without local changes to demography (Schaub and Ullrich 2021).

Critically, owing to the age of the samples we sequenced, our results are already more than a decade out of date. The most recently examined population of *cincta* from the Townsville Coastal Plain included in this study was sampled in November 2013. Black-throated finch flock sizes have declined in the Townsville Coastal Plain meta-population since 2006 (Mula Laguna et al., 2019) and some of the sites included in this study appear to have been locally extirpated within the last decade (SCG *pers. comm*.). Our findings highlight that the extirpation of any population results in the loss of unique genetic variation from the southern black-throated finch and will further increase this species’ risk of extinction. As recommended by the black-throated finch national recovery plan (BTFRT 2020), high-quality habitat must be identified, secured, and managed for conservation and connectivity. Moreover, the need for continued monitoring efforts and regular genetic surveys that enable measuring basic demographic parameters like immigration rate, relatedness structure, and reproductive success within both the Townsville Coastal Plain and the Galilee Basin meta-populations are essential for ensuring the long-term survival and recovery of the southern black-throated finch.

## Acknowledgements

The Australian National Wildlife Collection generously provided tissue and blood sample loans for black-throated finches, and we wish to thank Tony Grice, Stanley Tang, and Leo Joseph for their help with collecting and procuring samples used in this study. Illustrations of the black-throated finch and long-tailed finch were painted by Allison “AJ” Johnson. This project was funded by the Australian Research Council (ARC-DP-180101783 to DMH and SCG). DMH was supported by the Edward W. Rose Postdoctoral Fellowship, the Gerstner Scholars Fellowship and the Gerstner Family Foundation, and the Richard Gilder Graduate School at the American Museum of Natural History.

## Data Accessibility and Benefit-Sharing

Whole genome resequencing data for *Poephila cincta* and *P. acuticauda* are available in the Sequence Read Archive (BioProject PRJNA1147746 and PRJNA1101033) and European Nucleotide Archive (Project PRJEB10586). Reduced representation ddRAD data is available under SRA BioProject PRJNAXXXXXXX. This paper does not report original code, but our bioinformatics pipeline and custom analytical scripts have been deposited in GitHub, https://github.com/dhooper1/Black-throated-Finch. Any additional information required to reanalyze the data reported in this paper is available from the lead contact upon request.

## Author Contributions

DMH designed the research; DMH, KAL, and BB performed the research; DMH and KAL analyzed the data; SCG and IJL contributed critical resources; DMH wrote the paper with input from all authors.

---

**FIGURE S1.**
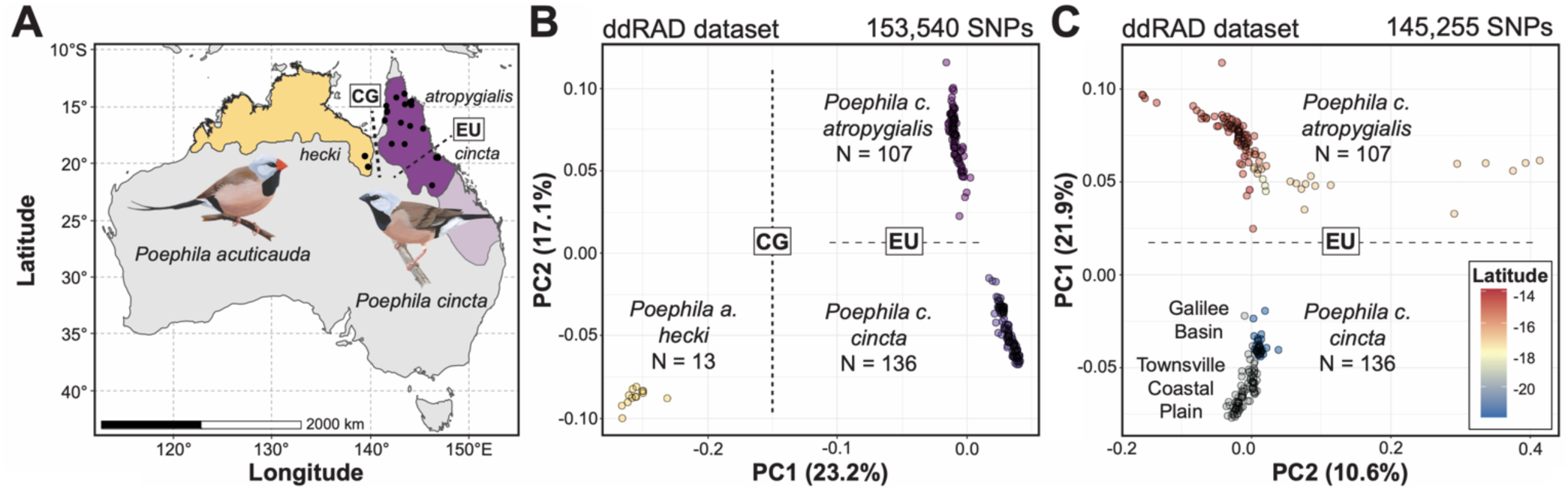
Geographic distribution and genomic divergence between the long-tailed finch (*Poephila acuticauda hecki*) and both black-throated finch (*Poephila cincta cincta* and *P. c. atropygialis*) subspecies. (A) Estimated range distribution of the long-tailed finch (yellow) and black-throated finch (purple). Recently extirpated regions of the black-throated finch geographic range (subspecies *cincta*) are represented with a lighter shade of purple. The geographic locations of populations sampled in this study are represented as black circles (see Table S3 for precise locations). The dashed lines represent the approximate geographic locations of two relevant biogeographic barriers: the Carpentarian Gap (CG) separating the long-tailed finch (west of barrier) and black-throated finch (east of barrier) and the Einasleigh Uplands (EU) separating black-throated finch subspecies *atropygialis* (north of barrier) from *cincta* (south of barrier). (B) Principal components analysis based on SNP variation using all ddRAD dataset samples from both species (N = 256) and (C) only samples from the black-throated finch (N = 243). Biogeographic barriers CG and EU are depicted as labeled dashed lines separating samples from either side of them. Samples are color-coded by taxon in (B) and by the latitude (i.e., LAT) of the population they were sourced from in (C). See Figure 1 for results with WGS dataset.

**FIGURE S2.**
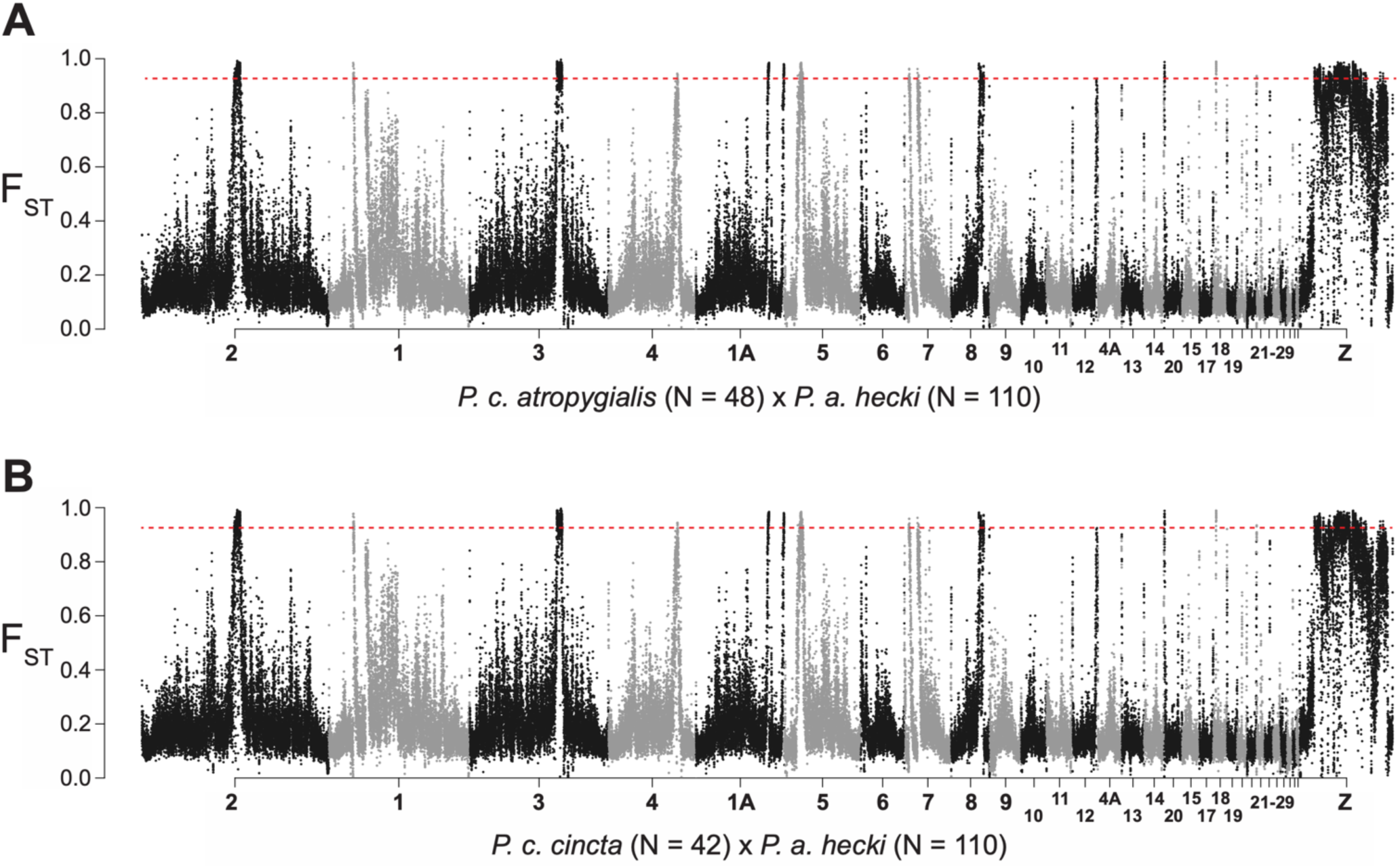
Genomic differentiation (F_ST_) between the long-tailed finch (N = 110) and black-throated finch subspecies (A) *atropygialis* (N = 48) and (B) *cincta* (N = 42) in 20 kb windows with 10 kb steps. The dashed red horizontal line represents the 99^th^ percentile threshold for the window based F_ST_ distribution (*atr-hec*, F_ST_ = 0.925; *cin-hec*, F_ST_ = 0.924).

**FIGURE S3.**
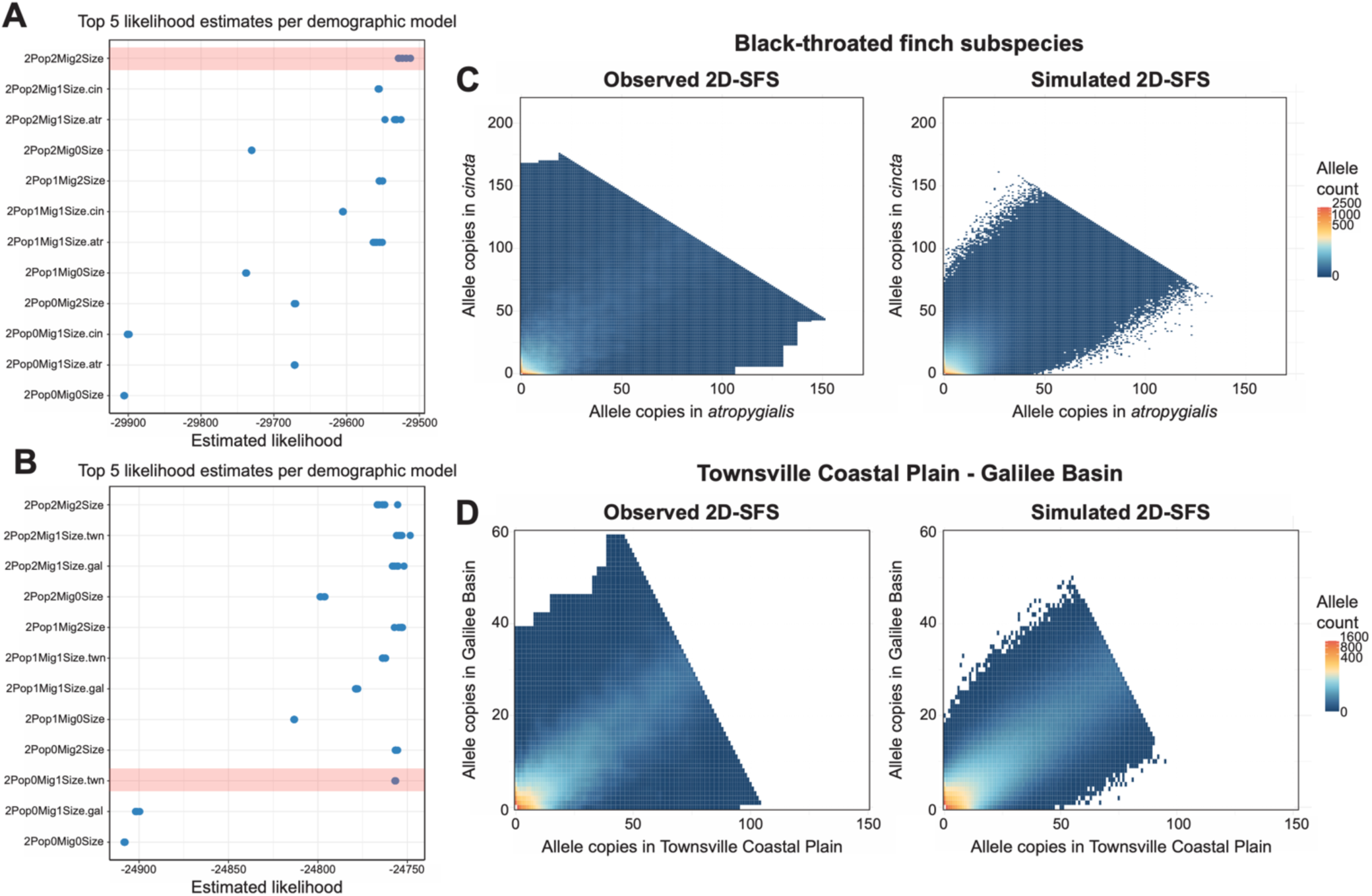
Demographic model inference. (A and B) Top five maximum likelihood scores plotted for 12 competing models, extracted from a set of 100 independent parameter searches per model. The red highlighted box indicates the best-fitting model based on the Akaike Information Criterion. Comparisons of the observed and simulated two-dimensional site frequency spectrum (2D-SFS) under the top-scoring model for (C) black-throated finch subspecies *atropygialis* and *cincta* (i.e., 2Pop2Mig2Size) and (D) between populations of southern black-throated finch subspecies *cincta* from the Townsville Coastal Plain and the Galilee Basin (2Pop0Mig1Size.twn). See Table S13 and Table S14 for parameter estimates under the best-fitting iteration of each model.

